# Environmental biofilms versus POCIS: efficiency to highlight environmental presence of pharmaceuticals and their effect on biofilm microbiome

**DOI:** 10.1101/2024.01.03.573948

**Authors:** Teofana Chonova, Agnès Bouchez, Leslie Mondamert, Elodie Aubertheau, Jérôme Labanowski

**Affiliations:** UMR CARRTEL, INRAE, USMB, F-74200 Thonon-les-Bains, France; Eawag, Swiss Federal Institute of Aquatic Science and Technology, 8600 Dübendorf, Switzerland; UMR IC2MP 7285, CNRS, Université de Poitiers, ENSIP, Poitiers, France

**Keywords:** environmental contaminants, passive samplers, WWTP effluents, river, diatoms, bacteria, DNA metabarcoding

## Abstract

Pharmaceutical compounds (PhC) are an important environmental issue, because of their high variety, potentially toxic byproducts and bioactivity at low concentrations. PhC concentrations in WWTP effluents often exhibit large and rapid variations that are difficult to record. Passive samplers are helpful to incorporate spot pollution events and register PhC occurrence at low concentrations.

In this work, we aim at (i) studying PhC accumulation in polar organic chemical integrative samplers (POCIS) and environmental biofilms exposed to urban (U) and hospital (H) treated effluents and (ii) evaluating the capacity of POCIS to predict changes in biofilm microbiome over a defined time period. Moreover, we (iii) determine the enrichment of PhC in the recipient river to evaluate levels of environmental contamination and potential effects on microbial biofilms. Biofilms and POCIS were installed in treated effluents and in the recipient river to measure the accumulation of PhC. In parallel, microbial biofilm communities were studied using DNA metabarcoding. The duration of each deployment was one month and the experiment was repeated six times. The performance of POCIS and biofilms to quantify PhC was depending on the compound. POCIS appeared well adapted to reveal contamination trends similar to these in the water column and to identify key PhC drivers of microbial changes. POCIS have the potential to predict pharmaceutical effects on biofilm community structure.

## Introduction

Wastewater treatment plants (WWTP) are one of the major sources of micropollutants for aquatic environments (Michael et al., 2013, Ternes and Joss, 2006). Their continuous discharges raise growing concerns about the health of these ecosystems (Schwarzenbach et al., 2006). In this context, hospital effluents are of great relevance. They usually contain higher varieties and concentrations of pharmaceutical compounds (PhC) that may persist during wastewater treatment and reach recipient aquatic environments (Acuña et al., 2020; Emmanuel et al., 2005; García-Galán et al., 2020; Heberer and Feldmann, 2005; Kosma et al., 2010; Rodriguez-Mozaz et al., 2020; Stadler et al., 2015; Wiest et al., 2018). PhC have become an important environmental issue, because of their high variety, potentially toxic byproducts and bioactivity at low concentrations (e.g. Ternes and Joss, 2006, Verlicchi et al., 2015). Designed to modify biochemical processes and to eliminate undesired microorganisms, they may affect non-target aquatic biota and may influence ecosystem processes that are mainly driven by microbial communities (Acuña et al., 2015; Corcoll et al., 2015; Proia et al., 2013b, 2013a).

Concentrations of PhC, especially in WWTP effluents, often exhibit large and rapid variations that are difficult to record with grab– and even with composite water sampling (Criquet et al., 2017). Representative water sampling can be difficult in physical, logistical and financial aspects (Alvarez et al., 2004). Passive samplers enable time-integrated measurement of PhC, incorporating daily and weekly variations and are therefore helpful to detect spot pollution events (Aubertheau et al., 2017; Criquet et al., 2017; Gravell et al., 2020; Guibal et al., 2018). Moreover, the accumulation of the target compound over time enables to record its occurrence at very low concentrations (Huerta et al., 2016; Miège et al., 2015). Thus, passive samplers may be helpful to record PhC presence in treated wastewaters and the resulting spot pollution events in the recipient environment. Different matrices can be used as passive samplers. Several studies have compared the efficiency of different man-made sampling devices to record pollutants and their correlation to water measurements (Criquet et al., 2017; Estoppey et al., 2019; Vermeirssen et al., 2012). Polar organic chemical integrative samplers (POCIS) are reliable and often used to record the presence of PhC (Camotti Bastos et al., 2018).

In aquatic environments, natural biofilms have the capacity to accumulate and transform a wide variety of contaminants including metals, pesticides and PhC (Achermann et al., 2019; Aubertheau et al., 2017; Bonnineau et al., 2020; Huerta et al., 2016; Mahler et al., 2020; Santos et al., 2019; Tien et al., 2013). This feature turns them into potential natural integrative samplers, useful for the detection of pollution hotspots (Aubertheau et al., 2017; Camotti Bastos et al., 2018; Fernandes et al., 2020; Rheinheimer dos Santos et al., 2020). Camotti Bastos et al. (2018) have shown that biofilms and POCIS can be used as complementary techniques for the identification of anthropogenic contamination. Moreover, environmental contamination is usually evaluated based on measurements in the water column and the transfer of pollutants to other environmental compartments is being neglected. Hence, complementary measurements in biofilms allow to better comprehend the transfer of pollutants and their potential impact on aquatic life. Biofilm microorganisms play an important role in the functioning of aquatic ecosystems (Battin et al., 2016). Several studies have shown negative effects of PhC on microbial biofilms (Corcoll et al., 2015; Proia et al., 2013b, 2013a; Rosi-Marshall et al., 2015, 2013). Hence, it is important to study the accumulation of PhC in biofilms to understand better microbial responses to PhC contamination and to preserve key ecosystem processes. Via measurements in the water column, Chonova et al. (2019, 2018a) showed that PhC are among the important factors structuring microbial biofilm communities in treated wastewaters and the recipient river. These findings opened a new question: To what extend do passive samplers have the potential to reveal the relationship between PhC concentrations and microbial community changes? Evaluating the response of microbial biofilms to PhC contamination as revealed by POCIS and biofilms will bring us a step further in the application of passive samplers for water quality assessment. However, the current state of art indicates a gap in understanding these relationships. To the knowledge of the authors, no previous environmental study has attempted to evaluate the accumulation of PhC in POCIS and biofilms for a controlled time period in order to study the potential of the integrative samplers to predict the response of microbial biofilms to contaminants.

In this context, our study aims at (i) investigating PhC accumulation in POCIS and environmental biofilms – exposed to urban (U) and hospital (H) treated effluents and (ii) evaluating the capacity of POCIS to predict changes in microbial biofilm communities over a defined time period. Moreover, we (iii) determine via POCIS the enrichment of PhC in the recipient river downstream from the WWTP output to evaluate levels of environmental contamination and potential effects on microbial biofilms.

## Materials and Methods

### Experimental design

The experimental site is a unique pilot WWTP that handles separately urban and hospital wastewaters using biological treatment with conventional activated sludge system and its recipient river (Labanowski et al., 2016; Chonova et al., 2018b) (Fig. S1A). This parallel separate treatment enables comparison of urban and hospital treated effluents (TE) and their effect on natural benthic microbial communities in the same socio-economical and geographical environment. Treated wastewaters are discharged into the recipient River Arve, which is used for drinking water production for Geneva (Switzerland).

The survey was performed in the urban (U) and hospital TEs (H) and the recipient river up-(RU) and downstream (RD) from the WWTP output. Stainless steel grid-baskets with autoclaved “blue schist” stones as substrates for natural biofilm colonization were installed in U and H and POCIS were installed in all four locations to collect in situ time-integrated PhC (Fig. S1C,D,E). The POCIS sampler consists of 200 mg (6cc) of resin as HLB sorbent with hydrophilic-lipophilic characteristics (Oasis®) compressed between two disk-shaped (90 mm diameter) microporous (0.1 μm) hydrophilic polyethersulfone (PES) membranes (Supor®). Stainless steel holdings were used to compress the membranes resulting in membrane open surface area of 22.9 cm^2^ that was protected from branches and rocks by stainless steel grid-baskets.

After each exposure period (1 month +/-2 days), POCIS were collected for the integrated measurement of PhC, the biofilm developed on the stones was scraped, suspended in sterile water and used to study the accumulation of PhC and the bacterial and diatom community composition. Stones were sterilized and reinstalled and new POCIS were submerged. All samples were transported to the laboratory in cool boxes close to 0° C for further analysis. This experiment was repeated six times from February to July 2014.

### Extraction and quantification of pharmaceuticals

A set of 18 PhC from 9 therapeutic classes were studied: Carbamazepine (CBZ), Caffeine (CAF), Propranolol (PROP), Levofloxacin (LVF), Iohexol (IOX), Diclofenac (DCF), Roxithromycine (ROXI), Ketoprofen (KETO), Sulfamethoxazole (SMX), 10,11-epoxy-carbamazepine (e-CBZ), Clofibrate (CLO), Atenolol (ATEN), Clofibric acid (ac-CLO), Trimethoprim (TMP), Bezafibrate (BZF), Metronidazole (MTN), Diazepam (DZP), Ranitidine (RND). Extraction from POCIS was performed on the AutoTrace^TM^ 150 Solid Phase Extraction (SPE) system with methanol as described in Bastos et al. (2018). Briefly, resins were transferred with ultrapure water to empty SPE cartridge (6 ml). The sorbent was washed with ultra-pure water, dried under gentle nitrogen flow, and weighted. Elution of POCIS was performed with 5mL of methanol (LC-MS/MS grade). The obtained extracts were evaporated under N_2_ flow until they reached a volume of 100 μL which was reconstituted in methanol/water (10:90, v/v). Extraction from biofilms were performed on an ASE™350 pressurized solvent extraction system at 80 °C, using methanol/water (1:2 in vol/vol) as extraction solvent. The extracts were then purified by SPE using Oasis® HLB cartridges (6 cc, 200 mg), with methanol as eluent. Final elution was carried out in the same way as for the POCIS.

PhC measurement was performed by liquid chromatography coupled to mass spectrometry (LC-MS) using a Q-Exactive Orbitrap (Thermo Fisher Scientific^TM^) and detected using an electrospray ion source operated in positive mode. Q-Exactive 2.0 SP 2 (application tune) software was used to control the mass spectrometer and the acquisition and data processing were carried out using Xashur 2.2 software, both Thermo Fisher Scientific (USA). The separation was done on an Acquity column UPLC® BEH C18 (2.1 x 100mm, 1.7 μm, Waters, Milford, USA) with methanol and water (both acidified with formic acid 0.3%) as mobile phase. The calibration was performed with a method of standard addition. The studied compounds were added in four concentrations (5, 10, 15, and 20 μgL^-1^) in the standard sample and they were injected three times with three replicates (Camotti Bastos et al., 2018; EU Norm NF ENV 13005, 1999). Limits of detection vary between 0.01 and 0.2 ng.g for POCIS and between 0.01 and 1 ng.g for biofilms. To study PhC measured in the water column, flow-proportional 24-h sampling was performed in U and H once a month on weekdays to measure a set of over 130 parameters listed in Chonova et al. (2018b) that included six of the molecules (ATEN, CBZ, DCF, KETO, PROP and SMX) studied in passive samplers here. PhC concentrations in the water column were analyzed in composite samples with high performance liquid chromatography coupled to a mass spectrometry (HPLC–MS/MS) as described elsewhere (Wiest et al., 2018).

### Microbial biofilm communities

In parallel to the integrative measurements of PhC, diatom and bacterial communities colonized during the same time period on the biofilms colonized in U, H, RU and RD were studied and were used to evaluate the capacity of the two passive samplers to reveal a correlation between the presence of PhC and microbial community changes in treated effluents. Microbial biofilm communities were sampled and analysed as described previously (Chonova et al., 2019). Briefly, the total genomic DNA was extracted, PCR amplification was performed using the primer pairs Diat_rbcL_708F (Stoof-Leichsenring et al., 2012) and R3 (Bruder and Medlin, 2007) for diatoms and F563 (Claesson et al., 2010) and 907rM (Schauer et al., 2003) for bacteria. Genomic libraries were prepared and sequenced on a PGM Ion Torrent sequencer. Bioinformatic processing was performed using Mothur Software (Schloss et al., 2009) as described elsewhere (Bailet et al., 2020; Chonova et al., 2019) to obtain operational taxonomic units (OTUs). This allowed to give an indication about the exposure of aquatic organisms to the contaminants present in the system and to study the potential of POCIS and biofilms to explain the response of microbial biofilm communities.

### Data analysis

PhC concentrations measured in passive samplers were calculated in ng.g.day^-1^. Quantitative (normalized abundances) matrices of bacterial and diatom OTUs were prepared to analyse microbial biofilm communities. All statistical analyses were performed with R 3.6.3 software (R Core Team, 2020) and graphical representations with ggplot2 package (Wickham, 2009).

Quantification frequency of PhC was calculated for each compound. Concentrations of individual PhC measured in POCIS and biofilms were visualized in a point chart. Pearson correlation tests (p < 0.05) were performed to study the relation between PhC measurements in the water column and integrated measurements in POCIS and biofilm.

Enrichment Factor (EF) was calculated to evaluate the accumulation of PhC in POCIS in the recipient river. Measurements in RU were applied as background values and EF was calculated as proposed by Aubertheau et al. (2017):

EF = log_10_([compound]_RD_/[compound]_RU_)

where [compound]_RU_ is the concentration of the given PhC in RU location, and [compound]_RD_ – in RD location. Positive values indicate an increase, and negative values indicate a decrease in the PhC concentrations downstream from the WWTP output. EF for each PhC and sampling period was visualized in a heatmap colored from white (low EF) to red (high EF). For data analyses, values under the limit of quantification were replaced by 1/2 of the respective LoQ value for each molecule.

Procrustes (Peres-Neto and Jackson, 2001, p.) and redundancy analyses (RDA) were used to assess the relationship between biofilm microbiome from TE sites and PhC measured in the two passive samplers. To that aim, PhC measurements from POCIS and biofilms were handled as two separate chemical datasets. The biological dataset was prepared by joining normalized quantitative OTU matrices of bacteria and diatoms.

Two Procrustes were performed to compare the correlation between microbial changes and PhC recorded in POCIS and in biofilms. After centring and scaling to unit standard deviation, two nonmetric multidimensional scaling (NMDS) analyses based on Euclidean distance similarity matrices were performed on the two chemical datasets and one NMDS based on Bray Curtis distance similarity matrix was performed on the biological dataset. Each Procrustes was done on the first two dimensions of the biological NMDS and of one chemical NMDS. The configurations were scaled to equal dispersions and a symmetric version of the Procrustes statistic was computed. The non-randomness between the two configurations was tested (protest, p < 0.05, 999 permutations).

In addition, two RDA were performed to infer the relationship and compute the biological variability, explained by PhC measurements in POCIS and biofilms. Prior to analysis, the biological dataset was Hellinger-transformed (Legendre and Gallagher, 2001). To reduce multicollinearity in chemical datasets, an approach based on principal component analysis (PCA) was used. One PCA was performed on each chemical dataset and the first two axes of each PCA were extracted and used as synthetic variables representative for the variability of measured compounds. Each RDA was performed on the biological dataset and on the synthetic variables extracted from one PCA. The sampling month was included as a covariate to consider the effect of time dependence. The significance was tested with permutation tests (999 permutations, *p* < 0.05).

Following the same procedure, a third Procruste and third RDA analysis were performed on river samples to assess the relation between microbial communities and PhC measured in POCIS in the recipient river. Multivariate analyses were done using the vegan package (Oksanen et al., 2019).

Finally, alpha diversity (number of OTU and chao1) of biofilm microbiome was calculated for all sampling sites. Pearson correlation tests were performed to study the relation between alpha diversity and integrated total (sum of all measured) PhC in POCIS (for treated effluents and the recipient river) and biofilms (for treated effluents only).

## Results and discussion

### Accumulation of pharmaceuticals into passive samplers

Measured concentrations of individual PhC in the HLB POCIS sorbent were ranging from <LOQ to 1354 ng.g^-1^.day^-1^ (for IOX) for treated effluent samples and from <LOQ to 48 ng.g^-1^.day^-1^ (for CAF) for river samples. Eleven out of 18 compounds (ATEN, CAF, CBZ, e-CBZ, DCF, IOX, KETO, PROP, ROXI, SMX and TMP) were quantified in at least 70% of the samples. BZF was quantified in 50% of the samples (never in H), MTN and DZP were quantified in around 30% of the samples (never in the river). CLO, ac-CLO, LVF and RND were not quantified in any POCIS sample (Table 1).

**Table 1.** – Quantification frequency of pharmaceutical occurrence measured in POCIS in treated effluents (TE) and in the river and in biofilms in TE. NSAID – non-steroidal anti-inflammatory drugs. LogK_ow_ – octanol water partition coefficient (https://pubchem.ncbi.nlm.nih.gov/ and https://www.drugbank.ca/)

| Pharmaceutical | Abbreviation | Therapeutic class | logK<sub>ow</sub> | Quantification frequency % |  |  |
| --- | --- | --- | --- | --- | --- | --- |
|  |  |  |  | Biofilm (TE) | POCIS (TE) | POCIS (river) |
| Clofibric acid | acCLO | lipid-lowering | 2.6 | 25 | 0 | 0 |
| Atenolol | ATEN | beta-blocker | 0.2 | 58 | 75 | 70 |
| Bezafibrate | BZF | lipid-lowering | 3.8 | 0 | 50 | 50 |
| Caffeine | CAF | stimulant | -0.1 | 100 | 92 | 100 |
| Carbamazepine | CBZ | anticonvulsant | 2.5 | 75 | 100 | 100 |
| Clofibrate | CLO | lipid-lowering | 3.3 | 8 | 0 | 0 |
| Diazepam | DZP | anxiolytic | 3 | 25 | 50 | 0 |
| Diclofenac | DCF | NSAID | 4.4 | 75 | 100 | 100 |
| 10,11-epoxy-carbamazepine | eCBZ | anticonvulsant | 1.3 | 17 | 100 | 50 |
| Iohexol | IOX | radiocontrast agent | -3.0 | 50 | 100 | 80 |
| Ketoprofen | KETO | NSAID | 3.1 | 83 | 100 | 80 |
| Levofloxacin | LVF | antibiotic | -0.4 | 100 | 0 | 0 |
| Metronidazole | MTN | antibiotic | 0.0 | 0 | 58 | 0 |
| Propranolol | PROP | beta-blocker | 3.0 | 100 | 100 | 70 |
| Ranitidine | RND | anti-acid | 0.3 | 67 | 0 | 0 |
| Roxithromycin | ROXI | Antibiotic | 3.1 | 83 | 92 | 50 |
| Sulfamethoxazole | SMX | Antibiotic | 0.9 | 67 | 100 | 80 |
| Trimethoprim | TMP | Antibiotic | 0.9 | 42 | 83 | 60 |

In biofilms, concentrations of individual PhC in TEs ranged from <LOQ to 45 ng.g^-1^.day^-1^ (for LVF). Six compounds (CBZ, DCF, KETO, ROXI, CAF, LVF and PROP) were quantified in more than 70% of treated effluent samples. IOX, e-CBZ and CLO were quantified in H only and BZF and MTN were not quantified in any biofilm sample (Table 1). CLO, ac-CLO, LVF and RND that were not detected via POCIS were recorded in biofilm samples. Aubertheau et al. (2017) have also quantified these four compounds in river biofilms.

### Major differences observed between POCIS and biofilms

PhC concentrations in POCIS were higher compared to biofilms. Such trend was reported previously also by Camotti Bastos et al. (2018). Different contaminants dominated POCIS and biofilm samples. LVF, CAF and PROP were the highest concentrated compounds in biofilms and IOX, CBZ, DCF, eCBZ, KETO and SMX – in POCIS (Fig. 1A).

**Fig. 1.**
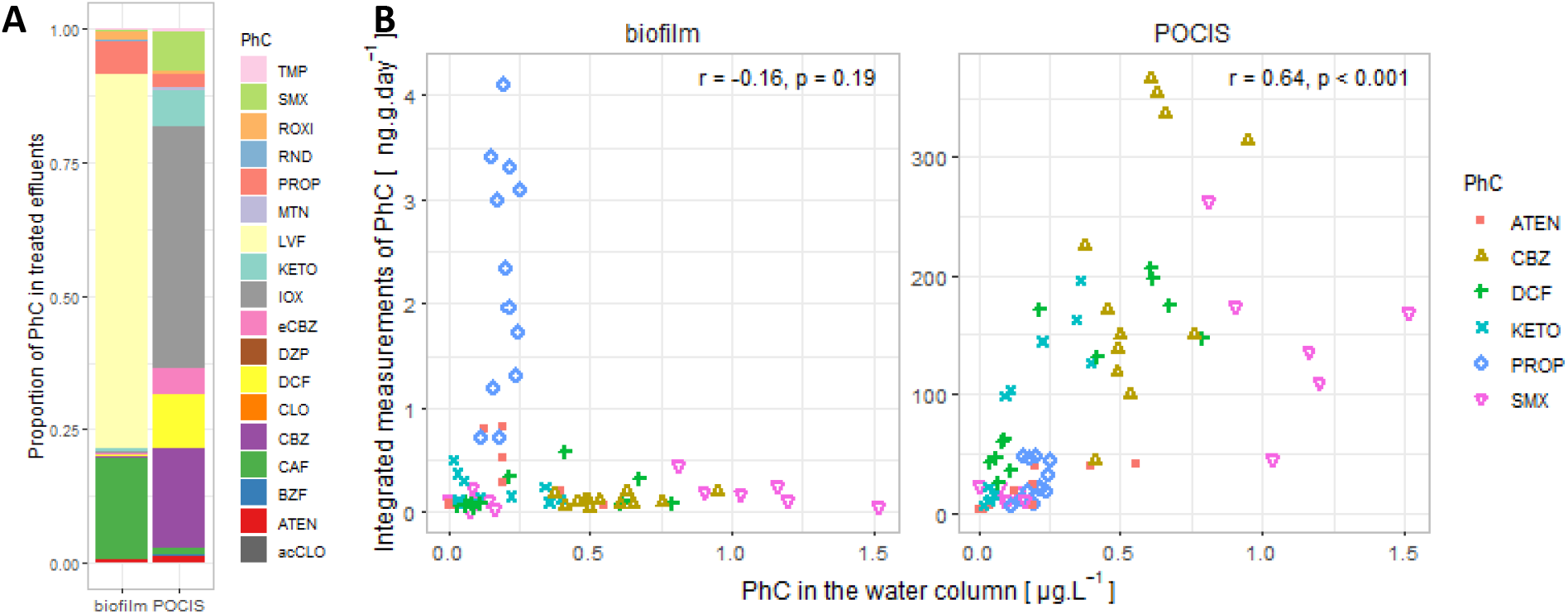
– (A) Stacked barplots presenting the proportion of PhC measured in biofilm and in POCIS in the treated effluents. (B) Correlations between PhC concentrations in the water column and in biofilms or POCIS. Pearson correlation coefficients (r) with the associated p-values are shown

Hydrophobic compounds with higher log*K*_ow_ values are likely to be retained in the PES membrane of POCIS, while highly polar or hydrophilic compounds with low log*K*_ow_ are expected to pass it more easily and to reach quickly the sorbent (Vermeirssen et al., 2012). Moreover, the affinity of the compound to the HLB sorbent depends on the functional groups of the molecule and the pKa (Yamamoto et al., 2009). Thus, some compounds are poorly retained on the HLB resin (Bäuerlein et al., 2012), while others bind well to the sorbent and are well detected in POCIS especially in concentrations close to the limit of detection. This was reported previously for TMP (Criquet et al., 2017). CBZ is also known for its high affinity to the HLB, because of its high number of sorption sites (Bäuerlein et al., 2012). Indeed, in this study, CBZ and TMP were detected at high frequency. In POCIS, the radiocontrast agent IOX was measured in highest concentrations compared to other contaminants (up to 1354 ng.g^-1^.day^-1^ in H). Given its polarity (log Kow = –3.0), IOX may easily pass the PES membrane and reach the sorbent (Vermeirssen et al., 2012).

As distinguished from POCIS, biofilms are natural components of aquatic ecosystems. Biofilms are mainly composed of bacteria, algae and fungi, embedded in a matrix of extracellular polymeric substances (EPS) and they are known as excellent indicators of pollution (Battin et al., 2016; Sabater et al., 2007). PhC can be accumulated both in the EPS and inside the microbial cells by active biological uptake.

Similarly to the findings of Aubertheau et al. (2017) and Huerta et al. (2016), in the present work we were able to detect in biofilms PhC with different charge and hydrophobicity (octanol-water partition coefficient logK_ow_ ranging from –3 to 4.4). However, dispite their successful quantification via POCIS, we, similarly to Aubertheau et al. (2017), failed to detect MTN and BZF in biofilms. Low concentrations of contaminants in the water column may also be a reason for this (Huerta et al., 2016). CBZ was detected in lower frequency in biofilm (75% of the samples) compared to POCIS (100%), which was also previously shown by Camotti Bastos et al. (2018).

### Correlation between pharmaceuticals in water and in passive samplers

Comparison of PhC measurements between composite water samples and integrated measurements showed significantly positive correlations (Pearson correlation tests, R = 0.64, p < 0.001) between water samples and POCIS, but not between water samples and biofilms (Fig. 1B). Strong positive correlations between composite water samples and POCIS were also reported previously (e.g. Criquet et al., 2017). By its design, the HLB resin exhibits a hydrophilic / lipophilic affinity balance, which allows the binding of polar compounds in addition to low-polar compounds (Vermeirssen et al., 2012). Due to its different components, the biofilm has a very large affinity potential, which allows to bind a wide range of pollutants but perhaps with competitive or modulating affinity effects (Flemming and Wingender, 2010). Moreover, PhC accumulated in biofilms may be less stable due to the biological processes driven by microorganisms (Tien et al., 2013; Winkler et al., 2001).

Daily and weekly fluctuations in PhC concentrations are often observed in WWTP effluents (Chonova et al., 2017; Goullé et al., 2012). The composite 24h water sampling enabled to incorporate daily variations, but not differences between working days and holidays. This may affect correlations between PhC concentrations measured in water and in passive samplers. Moreover, depending on their usage, some PhC may demonstrate short-term peak contamination, related to the treatment of disease outbreaks (Thomas et al., 2007). Such occurrences may be neglected or stronger represented in water samples. However, such behaviour is typical for molecules used for specific infections, but rather rare for largely used compounds like ATEN, PROP and TMP (Ort et al., 2010; Verlicchi and Zambello, 2016).

PhC are easily accumulated in environmental biofilms and sediments (Fernandes et al., 2020; Park et al., 2018). Still, environmental contamination of PhC is mainly evaluated via measurements in the water column and the transfer to other environmental compartments is regularly neglected. This may lead to regular underestimation of environmental contamination, especially for PhC that accumulate in biofilms and are therefore easily accessible to microorganisms (Oubekka et al., 2012). Such an example is PROP whose presence in the environment may be underestimated due to its strong affinity to the biofilm (Fig. 2B, Aubertheau et al., 2017). Our results highlight further the importance of reporting PhC concentrations in biofilms.

**Fig. 2.**
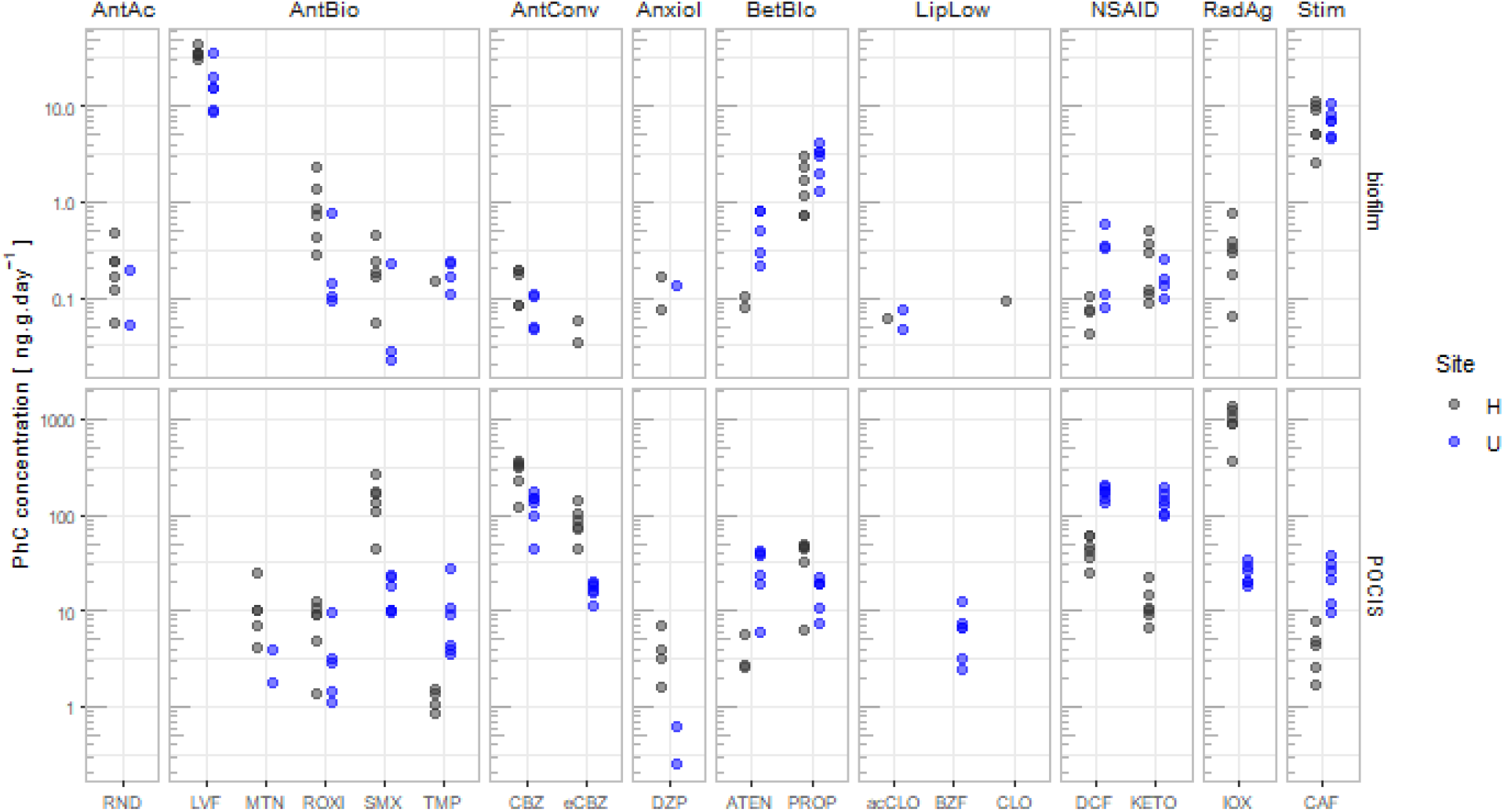
– Concentrations of individual PhC measured in biofilms and POCIS in urban (U, in blue) and hospital (H, in black) treated effluents (note the logarithmic scale). The compounds are grouped by therapeutic class (AntAc – anti-acid, AntBio – antibiotic, AntConv – anticonvulsant, Anxiol – anxiollitic, BetBlo – beta-blocker, LipLow – lipid-lowering, NSAID – non-steroidal anti-inflammatory drugs, RadAg – radiocontrast agent, Stim – stimulant). Measurements under the quantification limits are not shown

### POCIS and biofilms highlight pharmaceutical presence in urban and hospital WWTP effluents

Despite the different capacities of POCIS and biofilms to bind specific PhC, measurements in urban (U) and hospital (H) treated wastewater reveal that the two samplers indicate similar location-specific trends. This confirms the reliability of the two matrices to detect PhC contamination despite their specific advantages and shortcomings. Differences in PhC concentrations were observed when comparing U and H (Fig. 2, Fig. S2). Total concentrations were higher in H (1617±473 ng.g^-1^.day^-1^ measured in POCIS and 47±7 ng.g^-1^.day^-1^ measured in biofilms) followed by U (580±111 ng.g^-1^.day^-1^ measured in POCIS and 29±12 ng.g^-1^.day^-1^ measured in biofilms).

Contaminants detected in higher concentrations in H included CBZ, eCBZ, IOX, LVF (quantified in biofilms only), MTN (quantified in POCIS only), ROXI and SMX. As reported by POCIS measurements, IOX was quantified in all TE samples and it was around 40 times higher concentrated in H compared to U. In biofilm, IOX was quantified in H only. Such X-ray contrast media are usually given to patients in radiology departments and are excreted in hospitals. Clara et al. (2005) have also found previously that X-ray contrast media showed higher concentrations in wastewaters receiving hospital discharge, compared to exclusively urban wastewaters. However, depending on the need for hospitalization, around 30% of the patients may excrete such medicines outside hospitals (Kümmerer, 2004), which explains the presence of IOX in U (Kormos et al., 2011). As shown by both, POCIS and biofilms, all antibiotics (except for TMP) were detected in higher concentrations in H compared to U. The same trend was observed by measurements in the water column and it can be explained by the high and regular use of antibiotics in the hospital (Chonova et al., 2016). Antibiotics applied for hospital use often persist in the wastewater treatment and reach aquatic environments. This is a growing environmental and health concern, as the chronic exposure to antibiotics promotes the presence and the transfer of antibiotic resistance genes (Balcázar et al., 2015).

DCF, ATEN, BZF, KETO, TMP and CAF were generally detected in higher concentrations in U. Both matrices (POCIS and biofilms) reported higher concentrations of non-steroidal anti-inflammatory drugs (NSAIDs) in U compared to H (except for KETO in biofilms that did not show clear difference). This trend was also shown by measurements in the water column and it can be explained by the regular use of NSAIDs in domestic environments and their lower removal efficiency in the urban basin compared to the hospital one (Wiest et al., 2018).

CBZ, eCBZ, IOX and SMX exhibited highest concentrations in POCIS in H and DCF, KETO and CBZ in U (average exceeding 50 ng.g^-1^.day^-1^). Highest concentrations in biofilms were shown by LVF, PROP and CAF in both locations (average exceeding 1 ng.g^-1^.day^-1^).

### Transfer of pharmaceuticals to the recipient river

Time-integrated POCIS measurements enabled to incorporate potential spot pollution events that are likely to occur in a river downstream from a WWTP, but may be easily overlooked by water sampling (Criquet et al., 2017). In the recipient river, all measured PhC were found in higher concentrations downstream from the WWTP output compared to upstream, resulting in five times higher concentrations observed from RU (19±8 ng.g^-1^.day^-1^) to RD (105±41 ng.g^-1^.day^-1^) (Fig. 3A). In RU location, mean concentrations of individual compounds were below 1 ng.g^-1^.day^-1^, except for DCF and CBZ with 1.5 ng.g^-1^.day^-1^ and CAF with 13.4 ng.g^-1^.day^-1^. In RD location, DCF, CBZ, CAF, KETO and IOX exhibited mean concentrations above 9 ng.g^-1^.day^-1^. These results are in accordance with previous findings (Camotti Bastos et al., 2018). Enrichment Factor (EF) calculated to evaluate the accumulation of PhC discharged in the river reported solely postive values (ranging from 0.13 for CAF in July to 1.68 for PROP in June), indicating an increase downstream for all measured compounds during each sampling period (Fig. 3B). Molecules exhibiting highest EF were CBZ, eCBZ, DCF, KETO and PROP. The first four molecules were indeed among PhC measured in highest concentrations via POCIS in TEs (Fig. 2), which explains their high enrichment in the river.

**Fig. 3.**
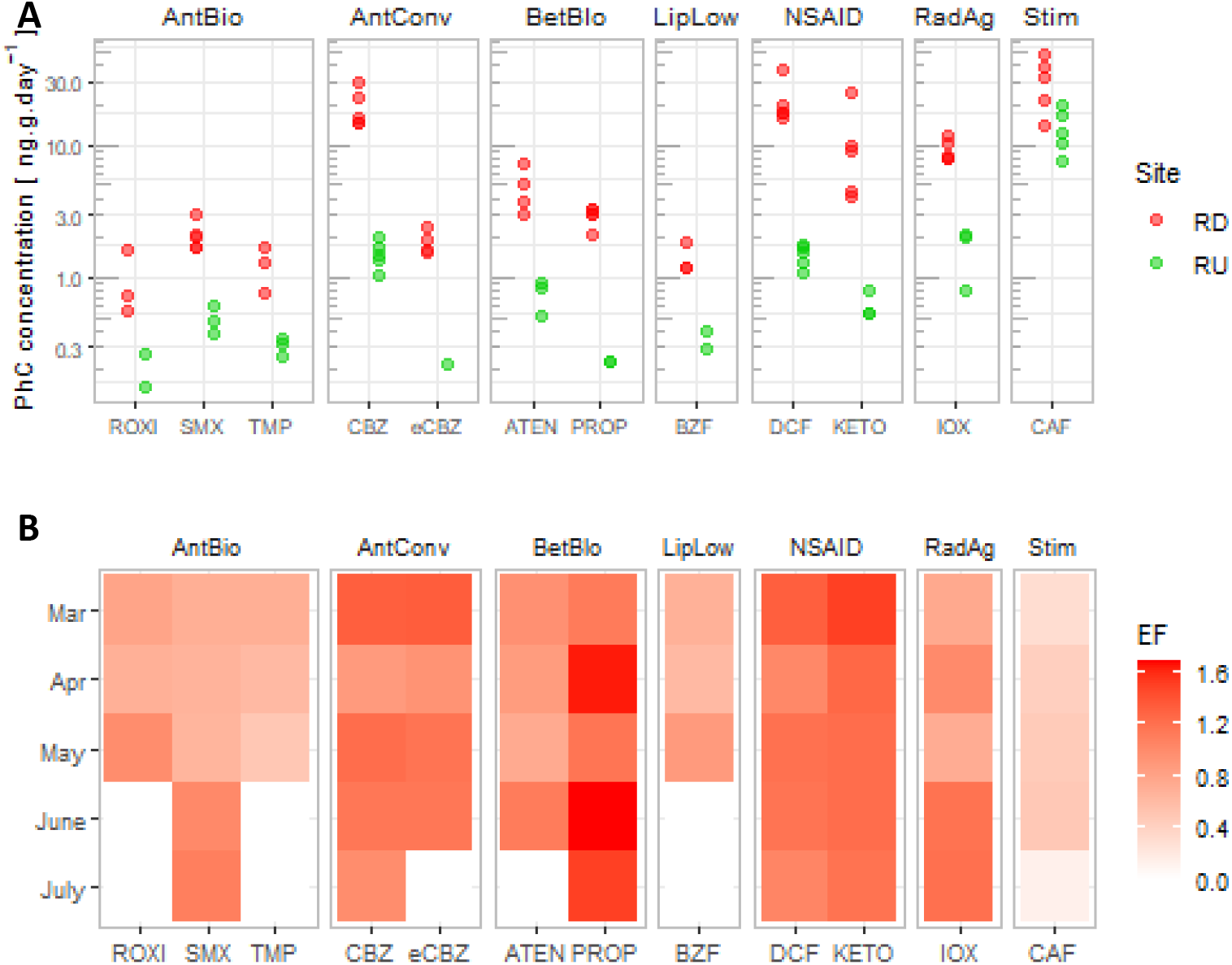
– (A) Concentrations of individual PhC measured in POCIS in the recipient river up-(RU, in green) and downstream (RD, in red) from the WWTP output (note the logarithmic scale). Measurements under the quantification limits are not shown. (B) Heatmap representing enrichment factor of PhC concentrations in RD with respect to RU for each compound and sampling period. White color represents measurements <LOQ. The compounds are grouped by therapeutic class (AntAc – anti-acid, AntBio – antibiotic, AntConv – anticonvulsant, Anxiol – anxiollitic, BetBlo – beta-blocker, LipLow – lipid-lowering, NSAID – non-steroidal anti-inflammatory drugs, RadAg – radiocontrast agent, Stim – stimulant)

Measurements <LOQ appeared mainly in warmer months (June and July). Temperature increase is expected to enhance sampling rates for POCIS (Greenwood et al., 2007; Ibrahim et al., 2013). The observed decrease of PhC concentrations in summer may be linked to degradation processes (Mathon et al., 2016; Yamamoto et al., 2009) or decrease in PhC consumption in summer resulting in lower concentrations that are difficult to detect (Chonova et al., 2017; Guibal et al., 2018; Huerta et al., 2016).

### Potential of passive samplers to reveal microbial changes related to pharmaceutical presence

Biofilms represent a majour part of the total microbial flora (up to 90%) in aquatic ecosystems and are responsible for vital functions including biogeochemical cycles and primary production (Battin et al., 2016). Numerous studies have reported negative effects of PhC on microbial biofilm structure and functioning. These are commonly expressed by decrease in biomass and diversity, decline in photosynthetic activity and bacterial enzymatic activity, increase in bacterial mortality rates, suppression of algal growth and microbial respiration and others (Corcoll et al., 2014; Drury et al., 2013; Eriksson et al., 2020; Guo et al., 2016; Proia et al., 2013b, 2013a; Rosi-Marshall et al., 2015, 2013; Serra-Compte et al., 2018; Tlili et al., 2020). It is therefore important to understand better the fate of these emerging contaminants in the different environmental compartments and their impact on aquatic organisms. It was previously shown via measurements in the water column that PhC are among the most important drivers of bacterial and diatom biofilm community changes in the treated wastewaters and in the river studied here (Chonova et al., 2019, 2018a, 2016). Integrated measurements of PhC are expected to give better insights on the impact of the effective PhC exposure over the complete biofilm colonization period. The application of DNA metabarcoding to study biofilm microbiome enables to record refined high-resolution genetic adaptations to these environmental pollutants.

Procrustes and RDA were used to study the efficiency of the two passive samplers to reveal relationships between PhC and genetic microbial changes. Procrustes found a significant correlation (protest, 999 permutations, p < 0.001) between PhC and biofilm microbiome in TEs for both passive samplers. RDA models were applied to investigate chemical effects on the microbiome taking into account the multivariate aspect of both – pharmaceutical measurements and microbial community structure. The two RDA that computed the biological variability explained by PhC measured with POCIS and biofilms, were significant (999 permutations, p < 0.05).

Most of the therapeutic classes studied here are known to affect microbial structure and functioning. Antibiotics are designed to directly target bacteria, which turns them into key drivers of these communities (Proia et al., 2013a). Changes in bacteria may indirectly affect algal composition. Beta-blockers, NSAIDs and analgesics were shown to inhibit bacterial development and algal photosynthesis (Bonnineau et al., 2010; Corcoll et al., 2014; Proia et al., 2013b). PhC are usually co-occurring in mixtures, whose effects raise concerns about the global environmental health (Bradley et al., 2016). The co-occurrence of a wide range of active compounds results in complex synergetic, additive or antagonist effects that are difficult to comprehend. Such cocktails of molecules were recently shown to disrupt key metabolic pathways responsible for biosynthesis of fatty acids and other important functions (Serra-Compte et al., 2018).

As revealed by Procrustes, PhC measured in POCIS showed higher correlation with microbial communities (r = 0.9, m^2^ = 0.18, Fig. 4B) compared to measurements in biofilms (r = 0.77, m^2^ = 0.41, Fig. 4A). The RDA model, revealing that POCIS explained 33% of the constrained variance and biofilms – only 29%, supported this finding (Fig. S3A, S4A,B). Hence, despite biofilms being the direct habitat of the studied microbiome, POCIS showed higher performance to reveal changes in their communities. This may be explained by the biological activities inside biofilms that cause transformation and biodegradation of contaminants by the microorganisms (Achermann et al., 2019; Ding et al., 2018, 2017; Santos et al., 2019; Tien et al., 2013; Winkler et al., 2001). Following these processes, bioavailable compounds, can be transformed to metabolites, conjugates and degradation products that may demonstrate toxic effects (Verlicchi et al., 2015). The presence of such byproducts cannot be recorded with the analytical methods used for mother compounds, thus we cannot account for their impact. Studying typical byproducts will help to better understand transformation processes occurring in biofilms and to better evaluate the potential impact on environmental microbiomes (Barbieri et al., 2012; Bonnineau et al., 2020). On the other hand, the EPS matrix of biofilms acts as a protection against contaminants and reduces their bio-accessibility and bioavailability to microorganisms. Oubekka et al. (Daddi Oubekka et al., 2012) suggests that the visco-elastic properties of extracellular substances may be a brake on the penetration of antibiotics. However, biofilm measurements of PhC include both, adsorbed (on the biofilm surface) and absorbed/internalized (in the EPS matrix/cells) compounds. Thus, measured concentrations involve molecules that may not have direct access and impact on biofilm microbiome. Analytical methods for separate measurements of biotic accumulation and abiotic sorption are necessary to better understand the microbial response to pollutants. However, these approaches are facing technical limitations so far (Bonnineau et al., 2020).

**Fig. 4.**
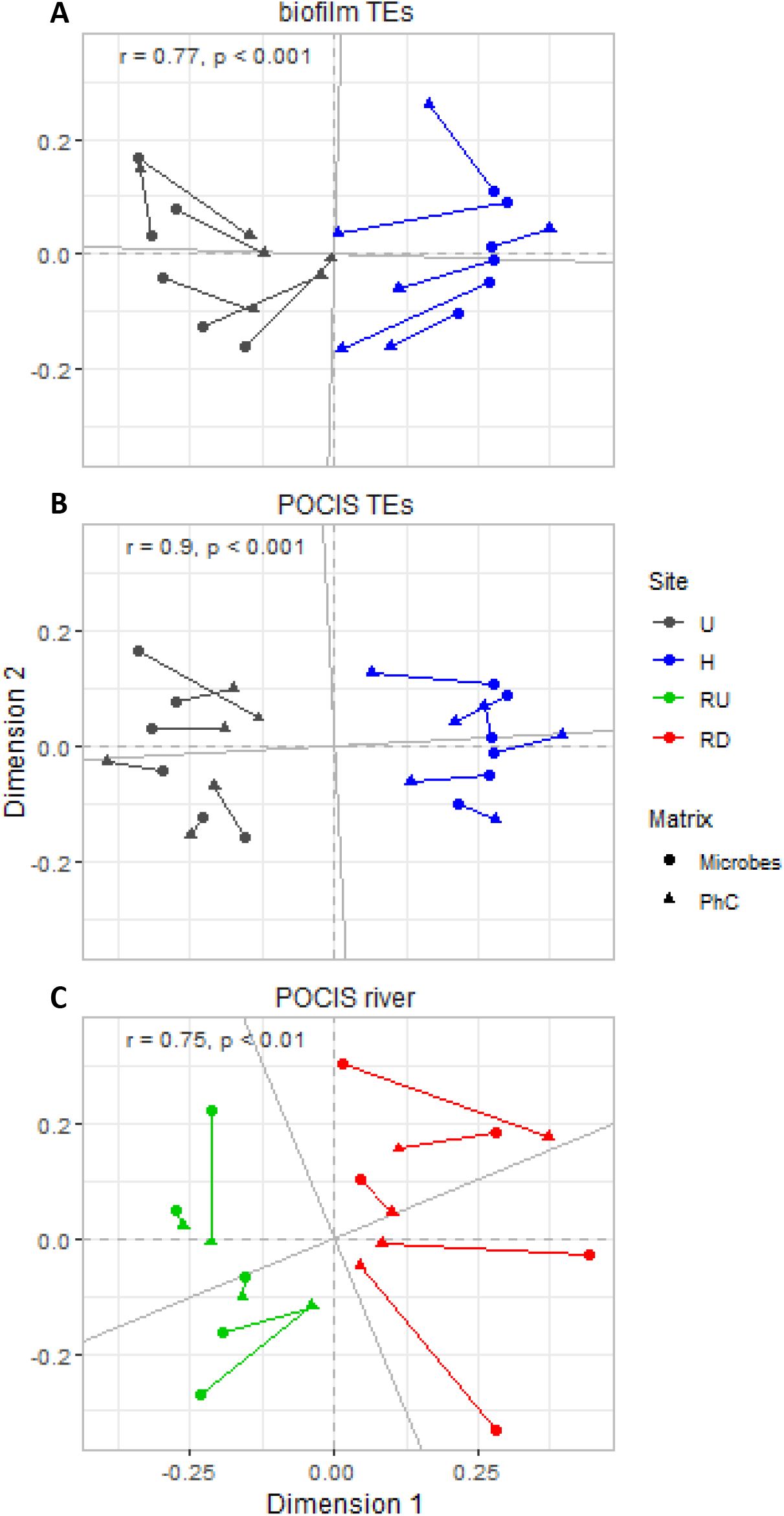
– Procrustes rotation presenting the relation between microbial biofilm community changes and concentrations of PhC measured in (A) biofilms and (B) POCIS, in urban (U, in blue) and hospital (H, in black) treated effluents (TEs) and in (C) POCIS in the recipient river up-(RU, in green) and downstream (RD, in red) from the WWTP output. Procrustes Correlations with associated p-values are shown

In the recipient river, significant correlation (Procrustes, r = 0.75, m^2^ = 0.43, p < 0.001, Fig. 4c) was also found between PhC concentrations recorded in POCIS and changes in biofilm microbiome. The RDA model accounted for 26% of the constrained variance, revealing the relation between river microbiome dynamics and the enrichment of PhC recorded in POCIS downstream from the WWTP output (Fig. S3B, S4C). Such microbial changes may include the replacement of taxa, sensitive to anthropogenic pressure and favor motile diatoms that are better adapted to pollution (Chonova et al., 2019; Rimet and Bouchez, 2011). The Procrustean correlation and the explained variance obtained for river samples were lower compared to TE samples. Indeed, rivers are complex ecosystems submitted to multiple pressures. Other factors like nutrient concentrations and water turbulences may also play a major role for the development and succession of microbial communities in river biofilms. Ecological guild classes of river benthic diatoms for example may show adaptations to micropollutants, but also to higher current velocity (Chonova et al., 2019; Passy, 2007).

Biofilm microbial alpha diversity was lower in treated effluents compared to the recipient river (Table S1). However, we could not find a significant correlation between alpha diversity (number of OTUs or Chao1) and integrated total PhC measured in POCIS and biofilms neither for treated effluents nor for the river locations. Hence, POCIS seem to be more effective to predict structural changes in microbial communities than changes in alpha diversity. Diversity indices reduce community data to one single number, discarding a great amount of information about important community characteristics (Biggs, 2000). This makes them less relevant to assess waterbodies pollution levels (Ricciardi et al., 2009; Pandey et al., 2017), which may explain the reduced potential of integrated PhC measurements to predict alpha diversity.

### Conclusions and perspectives for application in environmental monitoring

The robustness of the integrative sampling approach in combination with the cost-efficient ease of operation makes POCIS promising tools for regular water quality monitoring programs (Alvarez et al., 2004; Aubertheau et al., 2017; Camotti Bastos et al., 2018; Criquet et al., 2017; Harman et al., 2012; Poulier et al., 2014). However, the uptake processes of PhC in POCIS are complex and still poorly understood. Their uptake capacity is highly dependent on the characteristics of the aquatic environment (water flow and turbulences, temperature, etc.) and the molecule (log*K*_ow_, polarity, solubility, etc.) which makes it difficult to understand, model and predict PhC accumulation (Novic et al., 2017; Vermeirssen et al., 2012). Sampling rates (R_s_) for POCIS that are available in the literature to extrapolate measured concentrations in ng.g ^-1^ to ng.L^-1^ are often highly variable and based on calibrations in specific conditions that may not be representative for all aquatic environments (Ibrahim et al., 2013; Miège et al., 2015, Morin et al., 2012, Bailly et al., 2013). Thus, measures obtained from these integrative samplers, although promising, should still be interpreted with caution. Our results reveal that concentrations of PhC accumulated in POCIS resin in ng.g.day^-1^ appeared well adapted to reveal contamination trends significantly correlated to these measured in the water column and to predict chemical effects on biofilm microbial structure. POCIS benefit from a synthetic HLB sorbent matrix that facilitates the standardization and can bring an advantage over other natural compartments like sediments or biofilms. Contrary to POCIS, biofilms play a key role in the functioning of aquatic environments. They have a critical role in the repartition, transformation and removal of contaminants in the different compartments (Mahler et al., 2020). Biofilms are often used as bioindicators, because their microbiome, responsible for vital ecosystem functions, is highly sensitive to pollution. Structural and functional biofilm changes linked to the presence of PhC were previously reported (Aubertheau et al., 2017; Bonnineau et al., 2020; Huerta et al., 2016; Mahler et al., 2020; Santos et al., 2019). However, these processes are very complex and need to be better understood.

Our results showed poor correlations of biofilms with PhC measurements in other matrices and with microbial changes. This may be explained by the restrictive effect of the EPS matrix (Daddi Oubekka et al., 2012) and the transformation and degradation processes of contaminants (Achermann et al., 2019; Ding et al., 2018, 2017; Santos et al., 2019; Tien et al., 2013; Winkler et al., 2001). To better evaluate PhC presence and impacts in biofilms, byproducts should be included in future studies. Furthermore, improved analytical methods for separate measurements of compounds, adsorbed on the surface and such accumulated in the biofilm should be developed (Barbieri et al., 2012; Bonnineau et al., 2020). Despite improved technologies and analytical methods, the direct application of biofilms as standardized quantitative passive samplers in environmental biomonitoring will remain limited due to their biological nature. However, these improvements will help to better comprehend the transfer of contaminants from the water to biofilms and the following processes. The accumulation, transformation and effects of PhC in biofilms need to be studied further to better understand their impact on microbial and ecosystem functioning and their role in bioaccumulation and biomagnificantion processes that may affect not only aquatic microbiome, but also higher trophic levels (López-Doval et al., 2020; Ruhí et al., 2016, Bonnineau et al., 2020; Huerta et al., 2016).

## Abbreviations

10,11-epoxy-carbamazepine (e-CBZ), Atenolol (ATEN), Bezafibrate (BZF), Caffeine (CAF), Carbamazepine (CBZ), Clofibrate (CLO), Clofibric acid (ac-CLO), Diazepam (DZP), Diclofenac (DCF), Enrichment Factor (EF), extracellular polymeric substances (EPS), hospital treated effluent (H), Iohexol (IOX), Ketoprofen (KETO), Levofloxacin (LVF), Metronidazole (MTN), nonmetric multidimensional scaling (NMDS), pharmaceutical compound (PhC), polar organic chemical integrative sampler (POCIS), Propranolol (PROP), Ranitidine (RND), river downstream (RD), river upstream (RU), Roxithromycine (ROXI), Solid Phase Extraction (SPE), Sulfamethoxazole (SMX), treated effluents (TE), Trimethoprim (TMP), urban treated effluent (U), wastewater treatment plant (WWTP).

## Acknowledgements

We thank F.Rimet, R.Kurmayer, P.Illmer, F.Keck, L.Wiest, B.Gombert, A.LeBars and V.Vasselon for their contribution. We thank the Syndicat des eaux des Rocailles and Bellecombe for the permission to access and to experiment at the WWTP of Bellecombe.

## Consent to Publish

All authors have agreed with the content and all have given explicit consent to publish.

## Authors Contributions

Teofana Chonova, Agnès Bouchez, Leslie Mondamert, Elodie Aubertheau and Jérôme Labanowski contributed to the experimental design and sampling. Jérôme Labanowski, Leslie Mondamert and Elodie Aubertheau contributed to the chemical analyses. Teofana Chonova and Agnès Bouchez contributed to the molecular analyses. Teofana Chonova contributed to the bioinformatics, data organization and analysis, statistics, preparation of figures and writing of the manuscript. Jérôme Labanowski, Agnès Bouchez and Teofana Chonova contributed to the interpretation of the data and the revision of the manuscript. All authors approved the manuscript.

