## Supplementary material for "Environmental biofilms versus POCIS: efficiency to highlight environmental presence of pharmaceuticals and their effect on biofilm microbiome": Chonova_POCIS_SI.pdf

[2]Research Department for Limnology, Mondsee, Faculty of Biology, University of Innsbruck, Austria

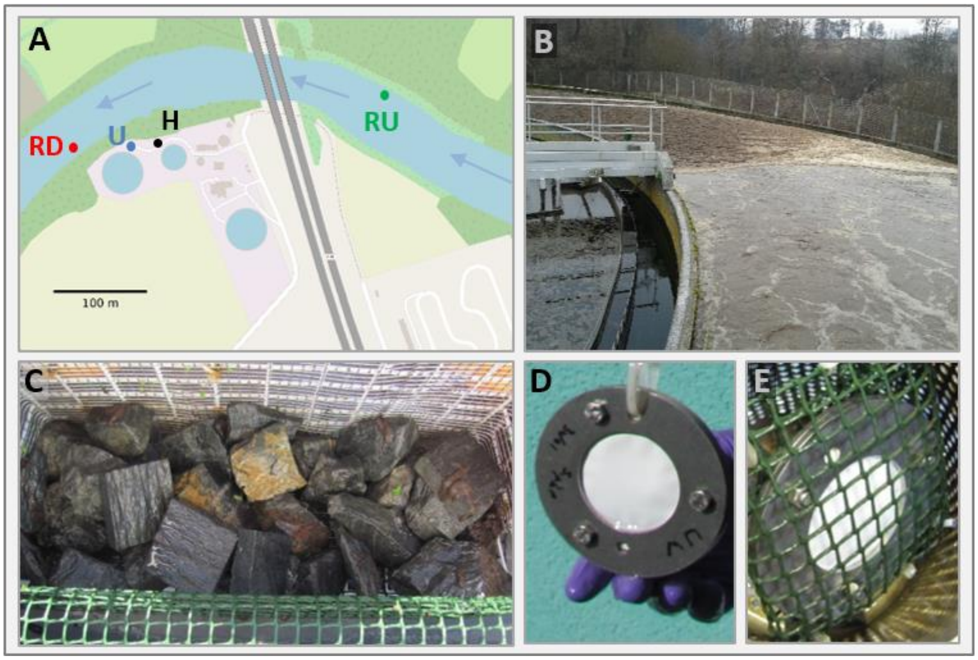


**Fig. S1** – (A) Sampling site map (U – urban treated effluent; H – hospital treated effluent; RU – river upstream; RD – river downstream; sampling locations are labelled with dots); (B) WWTP basin; (C) rocks installed for biofilm colonization; (D) polar organic chemical integrative samplers (POCIS); (E) POCIS in a protecting cage (Modified from Chonova et al., 2019)

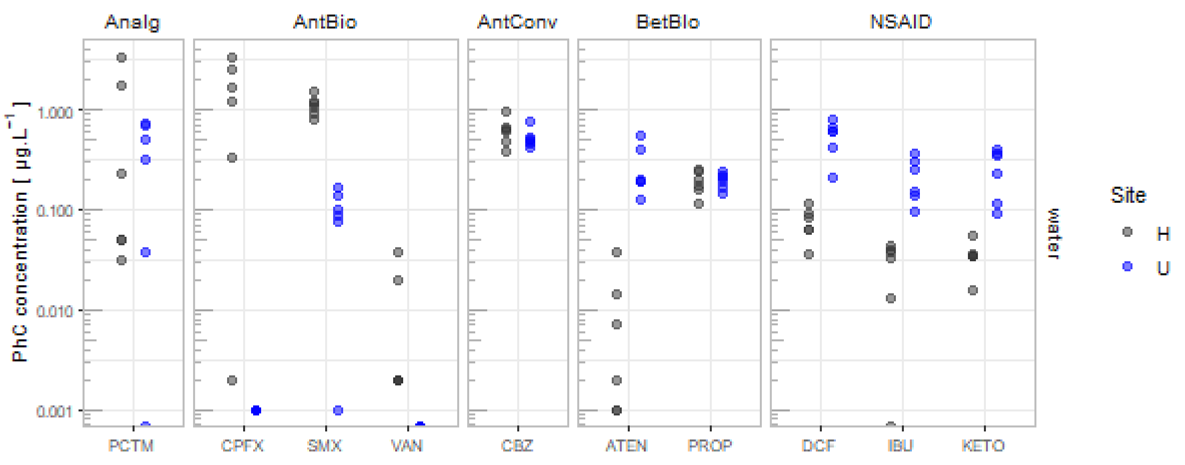


**Fig. S2** – Concentrations of individual PhC measured in the water column in urban (U, in blue) and hospital (H, in black) treated effluents (note the logarithmic scale). The compounds are grouped by therapeutic class (Analg – analgesics, AntBio – antibiotic, AntConv – anticonvulsant, BetBlo – beta-blocker, NSAID – non-steroidal anti-inflammatory drugs)

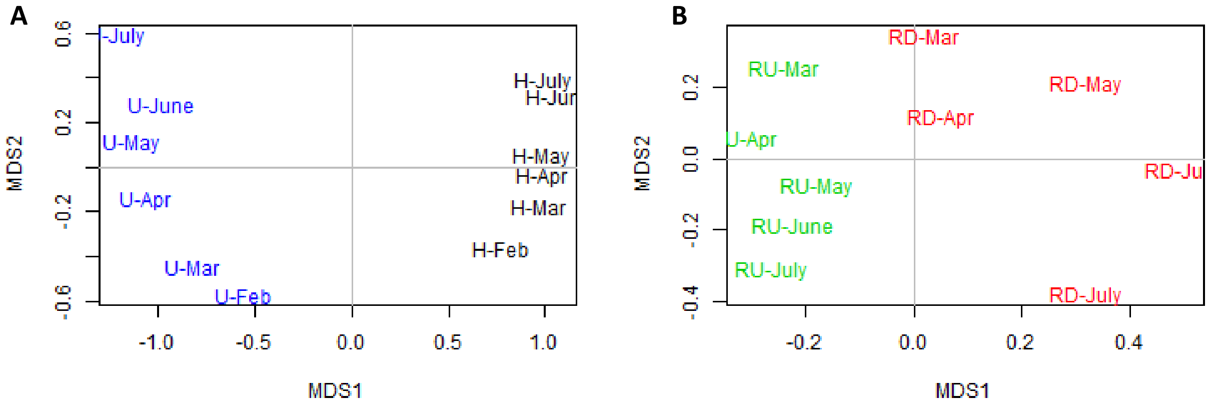


**Fig. S3** – NMDS two-dimensional plot of observed similarities between OTU profiles of microbial communities of (A) urban (U) and hospital (H) treated effluents and (B) river sites up- (RU) and downstream (RD) from the WWTP output

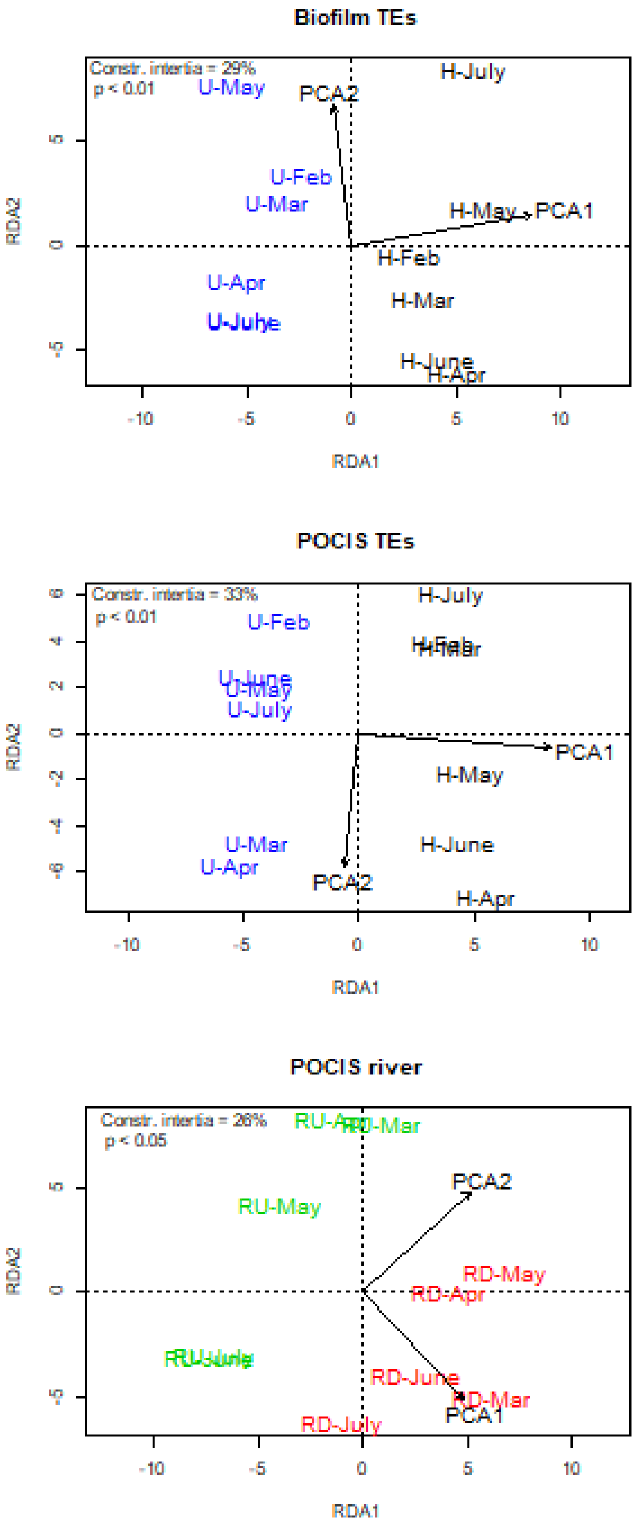


**Fig. S4** - Biplot from redundancy analysis (RDA) based on OTU profiles for periphytic samples and the variables PCA1 and PCA2 (synthetic variables corresponding to the first two PCA axes and representing the variability of measured compounds) from concentrations of PhC measured in (A) biofilms and (B) POCIS, in urban (U, in blue) and hospital

(H, in black) treated effluents (TEs) and in (C) POCIS in the recipient river up- (RU, in green) and downstream (RD, in red) from the WWTP output (Constr. Intertia = Constrained intertia, representing the percentage of explained variability in the biological dataset)

Table S1 – Number of OTU and chao1 calculated from OTUs of microbial communities developed in urban (U) and hospital (H) treated effluent sites and river sites up- (RU) and downstream (RD) from the WWTP output

| Sample | Number of OTU | chao1 |
|---|---|---|
| U-Feb | 844 | 919 |
| U-Mar | 980 | 1505 |
| U-Apr | 1027 | 1482 |
| U-May | 893 | 1464 |
| U-June | 1031 | 1603 |
| U-July | 847 | 1288 |
| H-Feb | 978 | 1034 |
| H-Mar | 1119 | 1820 |
| H-Apr | 1269 | 1811 |
| H-May | 1261 | 1950 |
| H-June | 1173 | 1687 |
| H-July | 1360 | 2048 |
| RU-Mar | 1360 | 2260 |
| RU-Apr | 1427 | 2506 |
| RU-May | 1766 | 2993 |
| RU-June | 1696 | 2838 |
| RU-July | 1605 | 2727 |
| RD-Mar | 1392 | 2472 |
| RD-Apr | 1633 | 2640 |
| RD-May | 1237 | 2116 |
| RD-June | 1273 | 2144 |
| RD-July | 1471 | 2771 |
